# Climate, crypsis and Gloger’s rule in a large family of tropical passerine birds (Furnariidae)

**DOI:** 10.1101/2020.04.08.032417

**Authors:** Rafael S. Marcondes, Jonathan A. Nations, Glenn F. Seeholzer, Robb T. Brumfield

## Abstract

Gloger’s rule predicts endothermic animals should have darker colors under warm and rainy climates, but empirical studies have typically found that animals tend to be darker under cool and rainy climates. Moreover, Gloger’s rule has rarely been tested jointly with the prediction that animals occupying dark habitats should have darker colors to enhance crypsis. We aimed to disentangle the effects of climate and light environments (habitat type) as correlates of plumage brightness in a large Neotropical passerine family. We found that cooler and rainier climates are associated with darker plumage, even after accounting for habitat types, and that darker habitats are associated with darker plumage, even after accounting climate. There was an important interaction between precipitation and temperature, whereby the negative effect of temperature on brightness becomes stronger under cooler temperatures. Climate and light environments have separate but complementary effects in driving macroevolutionary patterns of plumage color variation in birds.

---

Ecogeographic rules describe correlations between organismal phenotypes and features of their environment. Their repeated observation across taxa and space is *prima facie* evidence that they are driven by common selective pressures (Mayr 1963, James 1991, VanderWerf 2011). Gloger’s rule (Gloger 1833, Rensch 1929) is a longstanding ecogeographic rule describing a correlation between the colors of mammals and birds and the climatic conditions they occupy. Recent interpretations of Gloger’s rule (Delhey 2019, Marcondes et al. in review) have divided it into two versions: one “simple” and the other “complex”. The simple version relates to variation in overall melanin content, with greater amounts of melanin making feathers and fur darker (McGraw et al. 2005). This version of Gloger’s rule predicts that animals tend to be darker in rainy and warm climates and brighter in dry and cool climates (Gloger 1833). The complex version of Gloger’s rule concerns variation specifically in pheomelanin content (Delhey 2019), with greater amounts of pheomelanin making feathers and fur more brown or reddish-brown (McGraw et al. 2005). This paper concerns only the simple version of Gloger’s rule, which historically has been the only version most investigators have recognized (Delhey 2019).

Gloger’s rule has been investigated mostly at the intraspecific level, where evidence is abundant (Zink and Remsen 1986, Delhey 2019). Well-studied examples of species that have been found to be darker in more humid climates come from a broad variety of bird clades and include, but are not limited to, the Barn Owl *Tyto alba* (Roulin and Randin 2015, Romano et al. 2019), Black Sparrowhawk *Accipiter melanoleucos* (Tate and Amar 2017), Song Sparrow *Melospiza melodia* (Burtt and Ichida 2004) and Variable Antshrike *Thamnophilus caerulescens* (Marcondes et al. in review).

In contrast, Gloger’s (1833) prediction that animals should be darker in warmer climates has rarely been supported (Delhey 2019). More often, it has been found that populations inhabiting warmer climates tend to be lighter than their counterparts from cooler locales, a pattern dubbed Bogert’s rule and often attributed to thermoregulatory advantages (Clusella-Trullas et al. 2007, Rising et al. 2009, Delhey 2019).

Mayr (1956) argued that ecogeographic rules typically apply only to variation between populations within species, but Gloger’s rule has also been considered—and widely confirmed— at the interspecific level. In fact, interspecific comparative analyses are crucial to revealing how evolutionary processes operating within species can be generalized across macroevolutionary scales (Meiri 2011, Stoddard et al. 2019). The predicted negative correlation between brightness and humidity has been supported in phylogenetic comparative studies of the world’s primates (Kamilar and Bradley 2011), a large clade of Holartic shrews (Stanchak and Santana 2018), the entire Australian avifauna (Delhey 2018), the world’s woodpeckers (Miller et al. 2019), and the world’s passerine birds (Delhey et al. 2019). The latter two studies also supported the prediction of Bogert’s rule that animals are lighter in warmer regions.

Beyond climate, another major ecological axis to consider when investigating the causes of variation in animal color, particularly brightness, is habitat type, or light environment. Endler (1993) predicted that, to enhance crypsis, animals inhabiting dark light environments (*e.g*. the interior of dense forests) should be darker than those inhabiting open areas with bright light conditions (*e.g*., savannas), a prediction that has received wide support from comparative studies on birds (McNaught and Owens 2002, Gomez and Thery 2004, Dunn et al. 2015, Maia et al. 2016, Shultz and Burns 2017, Marcondes and Brumfield 2019). Because forests, particularly tropical rainforests, are more prevalent in rainier climates, this raises the possibility that the tendency for birds to be darker in more humid places (Gloger’s rule) is confounded by a need for crypsis in darker environments.

The passerine family Furnariidae (the woodcreepers, ovenbirds, foliage-gleaners and allies) is well-suited for investigating the relative roles of climate and light environments in driving interspecific variation in plumage brightness. Throughout the Neotropics, furnariids occupy virtually every terrestrial biome and habitat type (here construed to mean the spatial vegetation structure and density typically occupied by each bird species). They are found at the extremes of both precipitation and temperature in the Neotropics, from the warm and rainy Amazonian rainforests to warm and arid Chaco savannas, and from cool and dry high-elevation puna grasslands to the cool and rainy Andean cloud forests (Remsen 2003). Moreover, even under the same climatic conditions at a single geographic locality, species in this family specialize in such different habitat types as, for example, the lower strata of tropical rainforests, the forest canopy, and patches of open vegetation. They therefore experience dramatically different light environments, from the dim forest understory to intensely sun-lit fields and savannahs. Despite this ecological diversity, furnariids are virtually all festooned exclusively in innumerous shades of brown and rufous that vary relatively little in hue, but greatly in brightness. For example, furnariid colors range from light and creamy brown in the puna- and desert-inhabiting *Ochetorhynchus* earth-creepers to dark and rich brown in some species of tropical rainforest-dwelling *Xiphorhynchus* woodcreepers.

If Gloger’s rule is driven primarily by climate, species inhabiting rainy and warm climatic regimes are predicted to be darker than those from dry and cool regimes, regardless of their habitat preference. In contrast, if Gloger’s rule is mainly a result of birds adapting to be darker in darker (forest) habitats, bird species occupying forest habitats are predicted to be darker than their nonforest-based relatives, even if they inhabit similar climatic regimes. Marcondes and Brumfield (2019) previously demonstrated that furnariid species have evolved to be darker in darker habitats, consistent with Endler’s (1993) predictions for crypsis. Here, we sought to investigate the interacting roles of climate and habitat type in driving interspecific variation in plumage brightness in the Furnariidae.

## Methods

### Color data

We used the color dataset previously described in Marcondes and Brumfield (2019) and deposited on the Dryad digital repository under DOI 10.5061/dryad.s86434s (embargoed until July 16, 2021). Briefly, this dataset includes reflectance data for 250 (84%) furnariid species, with an average of 6.4 specimens per species (range: 1-8). For each specimen, this dataset includes reflectance spectra from seven plumage patches divided into a dorsal (crown, back, rump and tail) and a ventral (belly, breast and belly) set. We calculated plumage brightness (percentage of reflected white light) and averaged it across all specimens of each species. Because the Furnariidae are sexually monochromatic with no evidence of cryptic sexual dichromatism (Remsen 2003; Tobias et al. 2012; Diniz et al. 2016; Marcondes and Brumfield 2019), we considered the sexes together in our analyses. Finally, we used principal component analysis (PCA) to reduce our dataset to one principal component for the venter and one for the dorsum. The first principal components of dorsal and ventral PCAs were both loaded in the same direction by brightness in all plumage patches, thus representing overall brightness of that body surface; subsequent principal components captured various aspects of contrasts between plumage patches within each body surface (Marcondes and Brumfield 2019). Our final color dataset thus consisted of the first principal component score of brightness for each body surface (hereafter, simply “dorsal brightness” and “ventral brightness”) for each species.

### Habitat and climatic data

We used Marcondes and Brumfield’s (2019) categorization scheme for habitat types, which was based on Endler’s (1993) discussion of natural light environments. In brief, each of the 250 furnariid species we analyzed was assigned to one habitat type, in decreasing order of ambient light intensity: nonforest, intermediate and forest. The forest category includes only species that occupy the dimly-lit middle and lower strata of rainforests; we assigned canopy and edge species to the intermediate category because these areas are more intensely illuminated than the forest interior (Endler 1993, Marcondes and Brumfield 2019).

To obtain climatic data for each furnariid species we used the georeferenced locality records dataset of Seeholzer et al. (2017). This extensively-vetted dataset contains 23,588 occurrence records (average=70.4 records/species) gathered from museum specimens, audio recordings and observational records. For each locality in this dataset, we obtained mean annual temperature and mean annual precipitation from the BioClim database (Hijmans et al. 2005) and, for each species, we took the median of temperature and precipitation across all its occurrence localities. Because of their different magnitudes and units (°C for temperature and mm/year for precipitation), before fitting any statistical models (see below), at this stage we scaled each climatic variable to have a mean of 0 and a standard deviation of 1.

### Statistical analyses

To test the two hypotheses regarding the effects of climate and habitat on plumage brightness in the Furnariidae, we fit a series of phylogenetic Bayesian multilevel linear models using the modeling software Stan (Carpenter et al. 2017) as implemented in the R library *brms* (Bürkner 2017). All R scripts used for the analyses are available at https://github.com/jonnations. The multilevel model framework allowed us to fit linear models with multiple predictor variables while including group-level effects that account for statistical non-independence of species data due to shared phylogenetic history.

First, we tested the hypothesis that Gloger’s rule is primarily driven by climate, and species in wetter and warmer localities are darker than those from drier and cooler localities, regardless of habitat preference (forest, intermediate or nonforest). Under this hypothesis, we expect that species occupying wet and warm locales will be darker than those from dry and cool locales even when comparing nonforest species in the former to forest species in the latter. We fit two identical phylogenetic multivariate linear regression models, one with dorsal brightness as our response variable (Dorsal Model 1) and the other with ventral brightness as our response variable (Ventral Model 1). These models use precipitation, temperature, and the interaction between precipitation and temperature as the predictor variables. We used a species level matrix of scaled phylogenetic branch lengths (*i.e.*, the phylogenetic correlation matrix; Bürkner 2017) from the phylogeny of Harvey et al. (in review) as a group-level effect (de Villemereuil et al. 2012) to account for correlations due to phylogenetic relatedness of species. These models test the Gloger’s rule prediction that birds occupying warm and rainy regions should be darker than those occupying cool and dry regions (Gloger 1833, Rensch 1929). A nonzero, negative effect of precipitation on brightness would be consistent with Gloger’s rule, and a nonzero, positive effect would contradict it; likewise for the effect of temperature on brightness. Dorsal and Ventral Models 1 also estimate the interaction parameter of temperature and precipitation, which allows us to explicitly test whether wetter and warmer habitats result in darker plumage, and drier, cooler habitats result in brighter plumage. As both the dorsal and ventral brightness data had a slight positive skew, we used the skew-normal distribution family to describe the response variable rather than a simple Gaussian distribution. This distribution family estimates an additional parameter, alpha, which describes the direction and the strength of the skew. We fit regularizing priors on the group-level effects to prevent MCMC chains from occasionally searching very large, unreasonable values of model space (Gelman 2006; McElreath 2016). Each of the models included 4 chains run for 10000 generations with 5000 generations of warm-up and 5000 chains of sampling. We assessed chain convergence using the Gelman-Rubin diagnostic Ȓ, and chain efficiency using effective sample size (ESS). Ȓ < 1.01 and ESS > 500 represent acceptable convergence and mixing.

We also tested the alternative hypothesis that Gloger’s rule is related to light environments regardless of climatic variables. Under this hypothesis, we expect species occupying forested habitats to be darker than those inhabiting nonforest habitats, even if the nonforest species are in rainier and warmer climates. Specifically, we separated the effects of habitat from the effects of climate by fitting a phylogenetic multiple regression linear model with dorsal brightness (Dorsal Model 2) or ventral brightness (Ventral Model 2) as our response and temperature, precipitation, and our categorical habitat as predictor variables. As in Dorsal and Ventral Models 1, we included our phylogenetic correlation matrix as a group level effect. Dorsal and Ventral Models 2 have three predictor variables, generating three population-level outcomes: 1) The effect of precipitation on brightness, corrected for the influence of temperature, habitat, and phylogeny, 2) the effect of temperature on brightness, corrected for precipitation, habitat, and phylogeny, and 3) a posterior distribution of the mean brightness values of each habitat conditioned on the phylogenetic relationships, and corrected for the effects of precipitation and temperature. The estimated mean brightness values can directly address our question about whether species in darker habitats have darker plumage regardless of their climatic regimes. For categorical predictors, *brms* assigns a random category (habitat type in our case) as a dummy variable to use as the intercept value, so we removed the intercept parameter from the model to directly generate posterior distributions for each habitat. We used these posterior distributions of the mean brightness for each habitat to determine if species in different habitats differ in their brightness. To determine if the posterior distributions of the mean brightness for each habitat are different from one another, we calculated the distributions of the differences of each habitat’s brightness estimates, *i.e*. contrasts (nonforest-intermediate, nonforest-forest, intermediate-forest) (McElreath 2016; Roycroft et al. 2019) using the *compare levels* function in the R library *tidybayes* (Kay 2019). If the 95% credible interval of these difference distributions does not overlap zero, then we can credibly say that brightness is different between those habitats. This method of calculating the differences between posterior distributions is analogous to the “Bayesian T-Test” of Kruschke (2013). As in Model 1, we used regularizing priors and ran 4 chains of 10,000 generations and checked for convergence with Ȓ and ESS.

As a null model, we also fit an intercept-only phylogenetic multilevel model for brightness. This model has no predictor variables and only estimates the intercept of the group level-effect, in our case the phylogenetic correlation matrix. For each plumage surface (dorsal and ventral), we performed model comparison of our three models using the difference in expected log predictive density (ELPD) from the widely applicable information criteria (WAIC, Wantanabe 2010) using the *waic* function in the R package *loo* (Vehtari et al 2018), which calculates the ELPD and the standard error of the estimate. WAIC is appropriate for Bayesian inference with non-Gaussian posterior distributions (Gelman, Hwang, and Vehtari 2013); lower WAIC values represent greater support for a model. Comparing against the null allowed us to verify that precipitation, temperature, and habitat improved the predictive ability of our model rather than phylogeny alone explaining differences in brightness. WAIC also allowed us to assess whether the climate interaction model or the climate + habitat model was a better predictor of our brightness data.

## Results

All of our Bayesian multilevel models properly converged, and all parameters had Ȓ < 1.01 and ESS > 500. Results of Model 1 for the dorsal plumage showed a strong negative effect of precipitation on dorsal plumage brightness (Table 1, Figure 1), indicating that as precipitation increases, plumage gets darker, as predicted by Gloger’s rule. Model 1 also showed a strong positive effect of temperature on dorsal plumage (Table 1, Figure 1), demonstrating that as temperature increases, dorsal plumage gets brighter, *contra* Gloger’s rule but consistent with Bogert’s rule. We also found a positive interaction between precipitation and temperature (Table 1, Figures 1 and 2). This interaction indicates that the negative effect of precipitation on brightness decreases with increasing temperature. In other words, Gloger’s rule is more notable when comparing species that vary in the amount of precipitation they receive but all occupy similarly cool environments, rather than when comparing species that vary in precipitation but which all occupy similarly warm environments.

**Figure 1.**
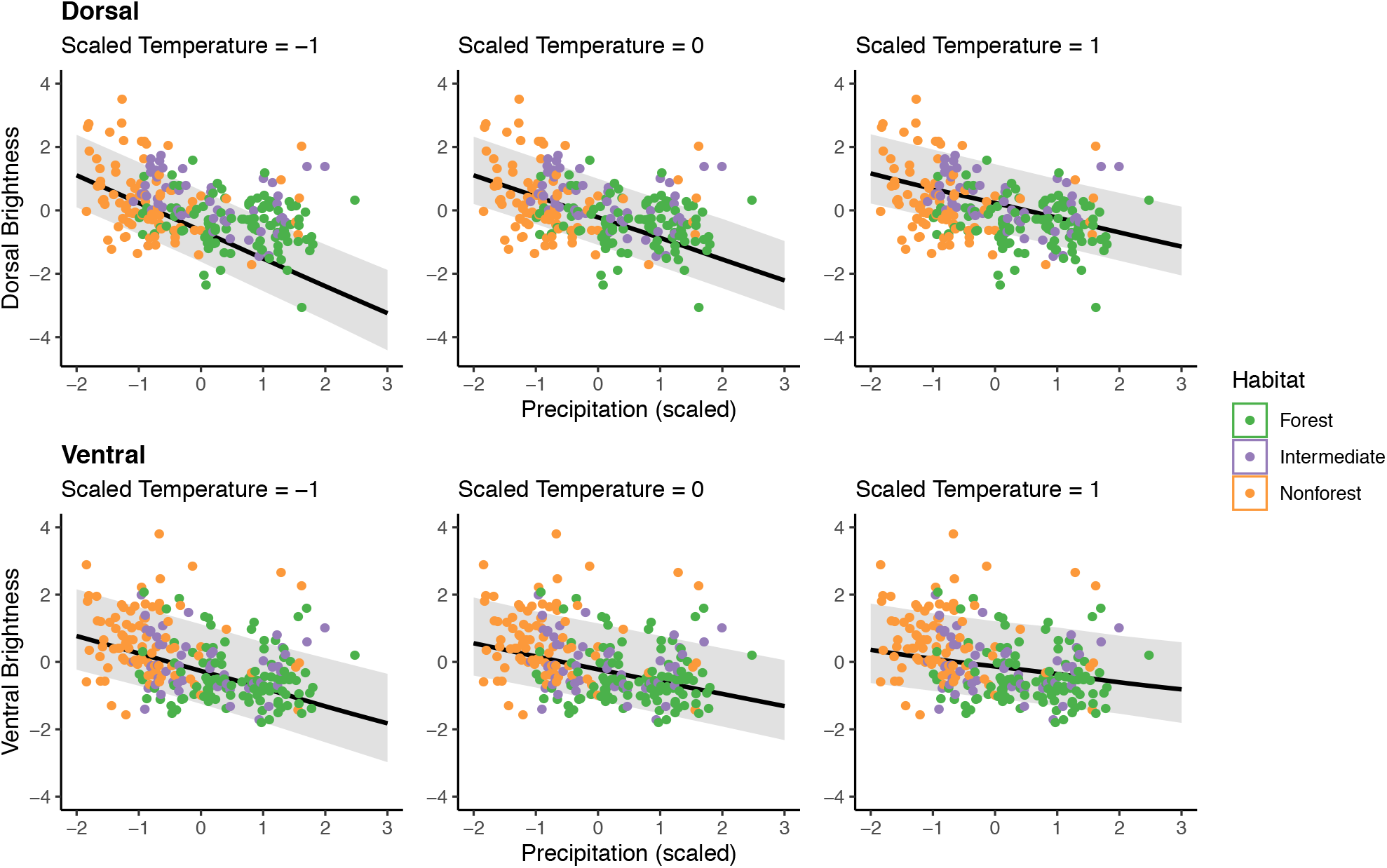
The slope of the negative relationship between temperature and brightness increases as temperature decreases (from Model 1). Plots in the left-hand column show the effect of precipitation on brightness when scaled temperature is −1 (mean – 1 standard deviation); the middle column shows the effect of precipitation on brightness when scaled temperature is zero (the mean), and the right-hand column shows the effect of precipitation on brightness when scaled temperature is 1 (mean + 1 SD).

**Table 1:**
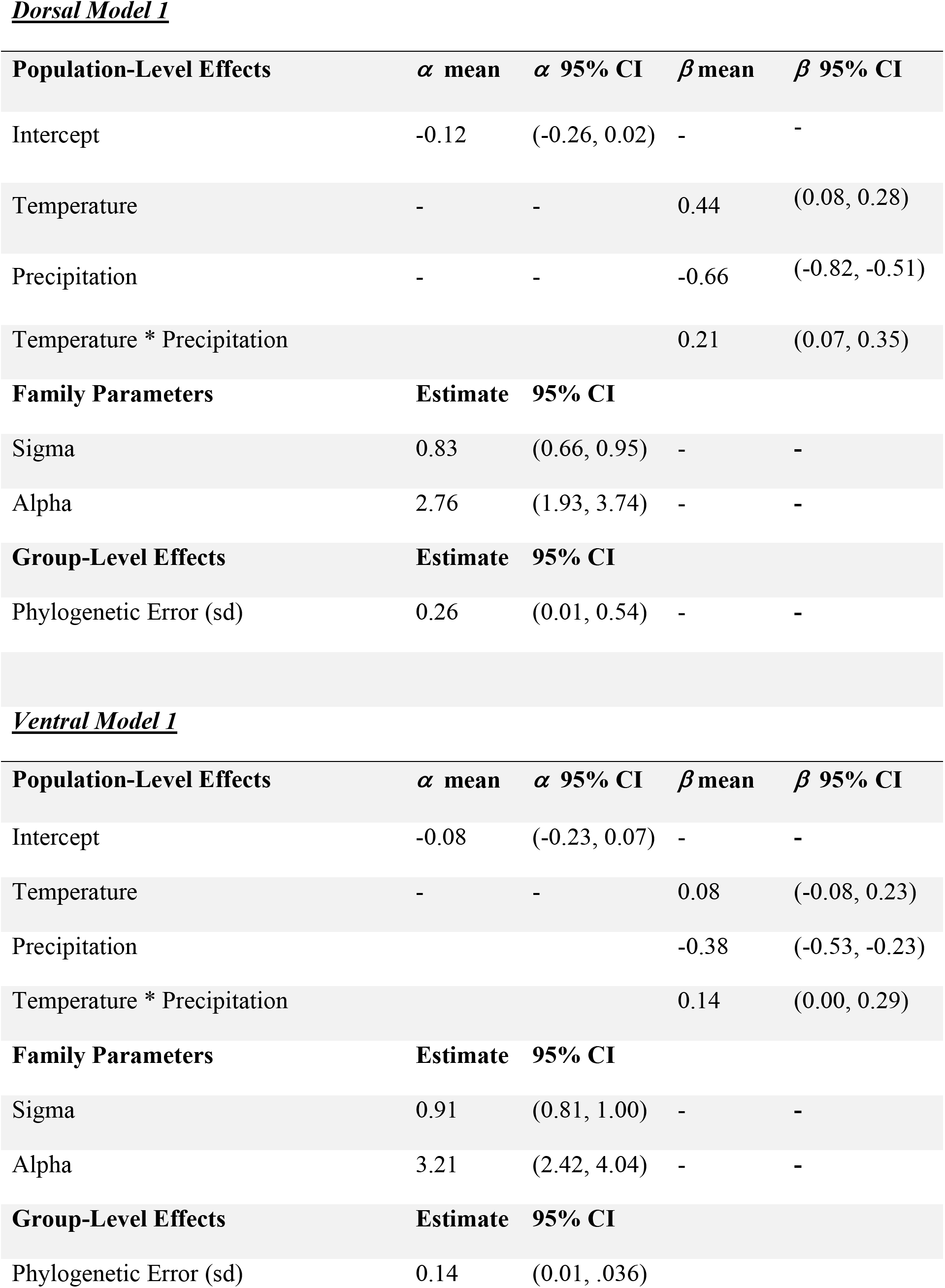
Results from Dorsal Model and Ventral Model 1, where plumage brightness is predicted by temperature, precipitation, and their interaction. *α* represents the intercept and *β* represents the regression coefficient conditioned on the phylogenetic correlation matrix. Population-Level Effects are the climatic parameters, Sigma is the residual error in the model, Alpha is the skew parameter in the skew-normal distribution, and Phylogenetic Error is the error in the model attributed to the phylogenetic correlation matrix.

**Dorsal Model 1**
| Population-Level Effects | $\alpha$ mean | $\alpha$ 95% CI | $\beta$ mean | $\beta$ 95% CI |
| --- | --- | --- | --- | --- |
| Intercept | -0.12 | (-0.26, 0.02) | - | - |
| Temperature | - | - | 0.44 | (0.08, 0.28) |
| Precipitation | - | - | -0.66 | (-0.82, -0.51) |
| Temperature * Precipitation |  |  | 0.21 | (0.07, 0.35) |
| Family Parameters | Estimate | 95% CI |  |  |
| Sigma | 0.83 | (0.66, 0.95) | - | - |
| Alpha | 2.76 | (1.93, 3.74) | - | - |
| Group-Level Effects | Estimate | 95% CI |  |  |
| Phylogenetic Error (sd) | 0.26 | (0.01, 0.54) | - | - |

Table 1: Results from Dorsal Model and Ventral Model 1, where plumage brightness is predicted by temperature, precipitation, and their interaction.
| Population-Level Effects | $\alpha$ mean | $\alpha$ 95% CI | $\beta$ mean | $\beta$ 95% CI |
| --- | --- | --- | --- | --- |
| Intercept | -0.08 | (-0.23, 0.07) | - | - |
| Temperature | - | - | 0.08 | (-0.08, 0.23) |
| Precipitation |  |  | -0.38 | (-0.53, -0.23) |
| Temperature * Precipitation |  |  | 0.14 | (0.00, 0.29) |
| Family Parameters | Estimate | 95% CI |  |  |
| Sigma | 0.91 | (0.81, 1.00) | - | - |
| Alpha | 3.21 | (2.42, 4.04) | - | - |
| Group-Level Effects | Estimate | 95% CI |  |  |
| Phylogenetic Error (sd) | 0.14 | (0.01, .036) |  |  |

We found similar results for the ventral plumage. Precipitation had a negative effect on ventral plumage brightness (Table 1, Figure 1). Temperature had an uncertain positive effect on ventral brightness (95% credible interval overlapping 0), demonstrating that temperature has less importance on ventral plumage than on dorsal plumage. We found a weak positive effect of the interaction between precipitation and temperature on the ventral plumage (Table 1, Figures 1 and 2).

There was a subtle difference between the venter and the dorsum in effects of the interaction of precipitation and temperature and brightness (Figure 2). Both plumage surfaces tended to be darkest for species in cool/rainy climates and brightest in cool/dry climates. But whereas temperature seemed to have little effect on dorsal brightness in dry climates, ventral plumages tended to be darker under warm/dry conditions than under cool/dry conditions. In other words, there is little change in dorsal brightness when comparing species from cool/dry and warm/dry conditions, but ventral brightness is higher (lighter) in species from cool/dry than warm/dry conditions.

**Figure 2.**
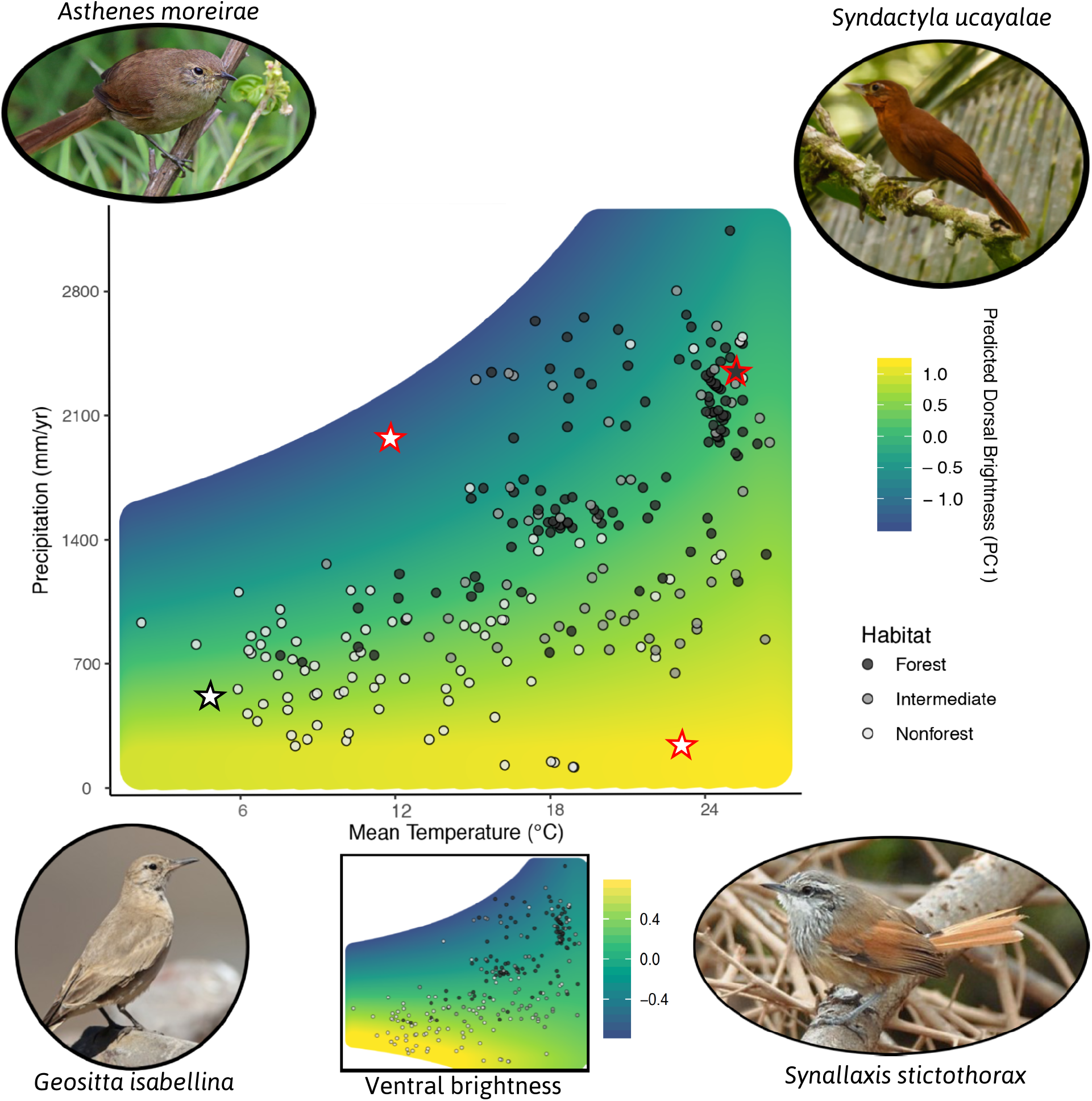
A complex interaction between precipitation and temperature predicts dorsal (main panel) and ventral (inset at bottom) brightness in the Furnariidae. The colors in the heatmap represent brightness as predicted by Model 1, which includes precipitation, temperature and their interaction as predictors. Stars represent the depicted species they are closest to. Photograph authors and licenses: Top left: Nigel Voaden CC BY-SA 2.0. Top right: Rubens Matsushita ©. Bottom right: Roger Ahlman ©. Bottom left: Luke Seltz ©.

Our Dorsal and Ventral Models 2, in which we removed the interaction between precipitation and temperature and added habitat as a predictor, showed a similar negative effect of precipitation on both dorsal plumage and ventral plumage (Table 2). We also found positive effects of temperature on dorsal, and an uncertain effect of temperature on ventral plumage (Table 2). This model also estimated the posterior distributions of mean plumage brightness for each habitat, conditioned on phylogenetic effects and the climatic variables (Figure 3). We then calculated the differences of those distributions (Figure 3). For the dorsal plumage we found that intermediate and nonforest species are credibly brighter than forest species but not distinguishable from each other, because the difference in their posterior distributions overlapped zero (Figure 3). Results were similar for ventral plumage, except that the difference between intermediate and forest species slightly overlapped zero. Therefore, we found that birds in forest habitats are darker than birds from intermediate or nonforest habitats, even after accounting for differences in temperature and precipitation.

**Figure 3.**
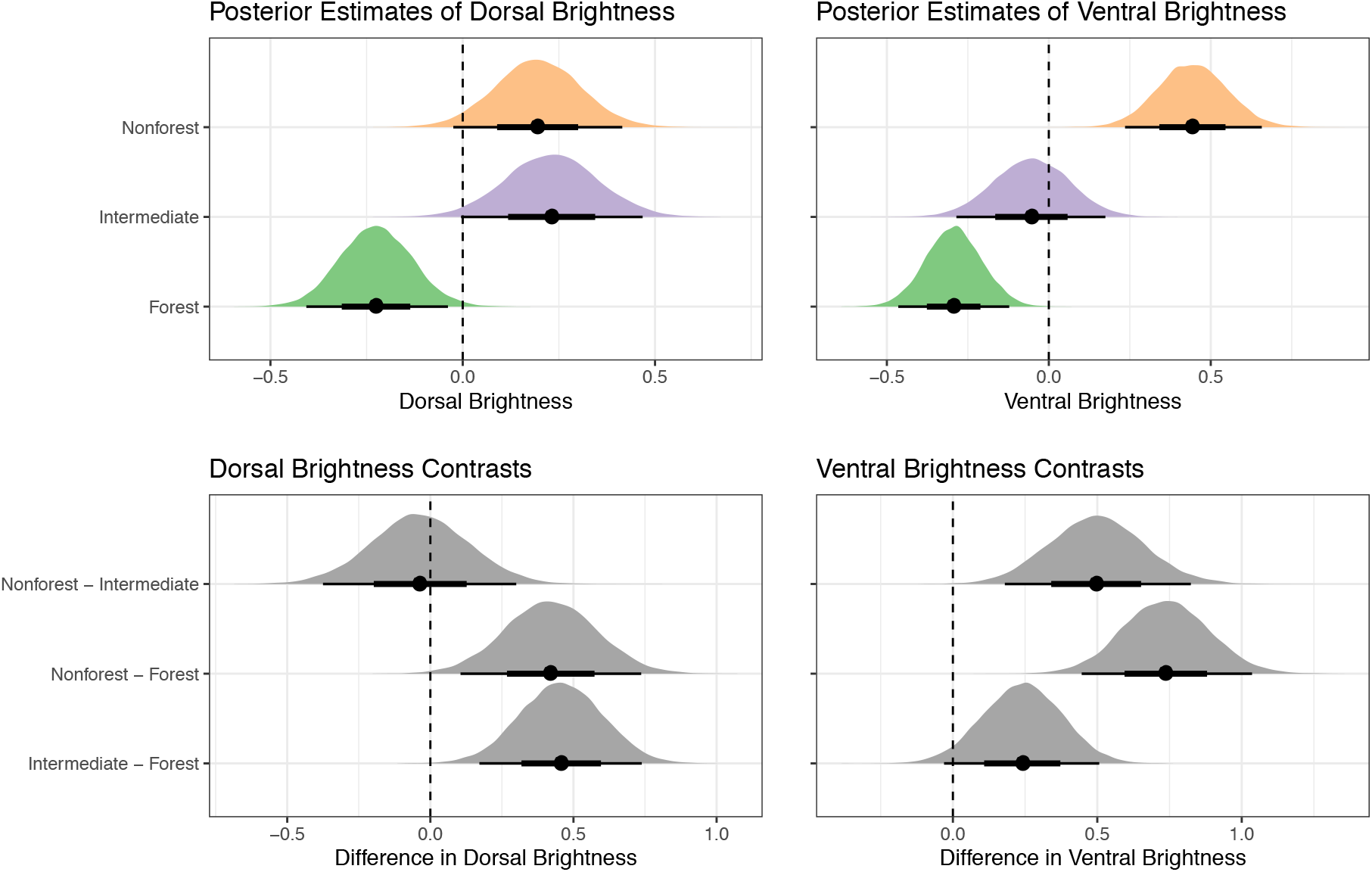
Differences in mean plumage brightness across light environments persist even after accounting for climatic variation. Top row: posterior distributions of the mean brightness value, conditioned on temperature, precipitation, and the phylogenetic correlation matrix, for the effects of climate. Bottom row: contrasts between the phylogenetic means of each habitat. If the contrast overlaps zero (dotted line), then there is no credible difference between the brightness of the two habitats. The black circle represents the mean and the horizontal bars the 66% (thick bar) and 95% (thin bar) credible intervals of the distribution.

**Table 2:**
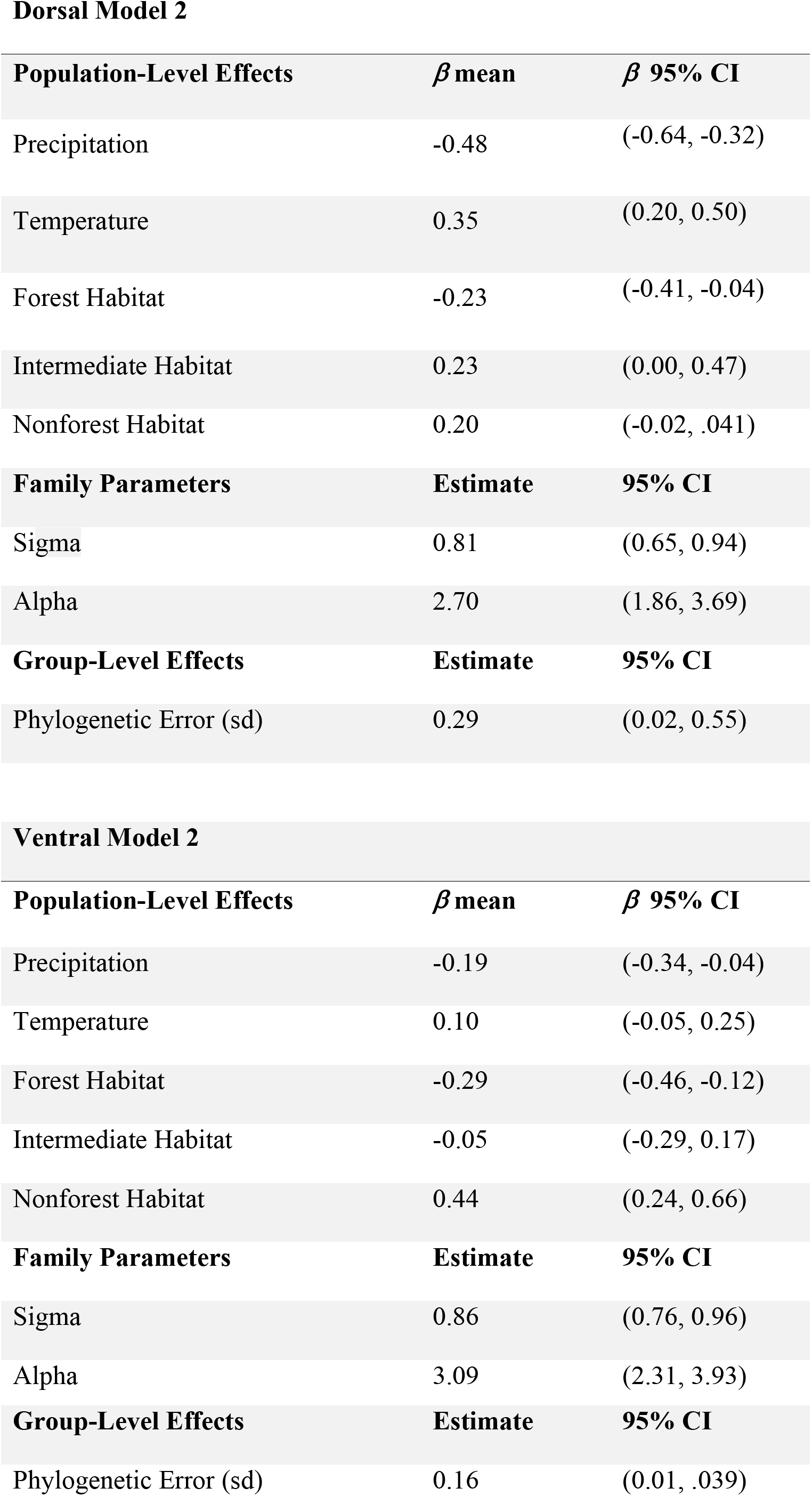
Results from Dorsal Model and Ventral Model 2, where brightness is predicted by temperature, precipitation and habitat type. *β* represents the regression coefficient, conditioned on the phylogenetic correlation matrix. Population-Level Effects are climatic parameters and the brightness estimates for each habitat. Sigma is the residual error in the model, Alpha is the skew parameter in the skew-normal distribution, and Phylogenetic Error is the error in the model attributed to the phylogenetic correlation matrix.

**Dorsal Model 2**
| **Population-Level Effects** | **$\beta$ mean** | **$\beta$ 95% CI** |
| --- | --- | --- |
| Precipitation | -0.48 | (-0.64, -0.32) |
| Temperature | 0.35 | (0.20, 0.50) |
| Forest Habitat | -0.23 | (-0.41, -0.04) |
| Intermediate Habitat | 0.23 | (0.00, 0.47) |
| Nonforest Habitat | 0.20 | (-0.02, .041) |
| **Family Parameters** | **Estimate** | **95% CI** |
| Sigma | 0.81 | (0.65, 0.94) |
| Alpha | 2.70 | (1.86, 3.69) |
| **Group-Level Effects** | **Estimate** | **95% CI** |
| Phylogenetic Error (sd) | 0.29 | (0.02, 0.55) |

**Ventral Model 2**
| **Population-Level Effects** | **$\beta$ mean** | **$\beta$ 95% CI** |
| --- | --- | --- |
| Precipitation | -0.19 | (-0.34, -0.04) |
| Temperature | 0.10 | (-0.05, 0.25) |
| Forest Habitat | -0.29 | (-0.46, -0.12) |
| Intermediate Habitat | -0.05 | (-0.29, 0.17) |
| Nonforest Habitat | 0.44 | (0.24, 0.66) |
| **Family Parameters** | **Estimate** | **95% CI** |
| Sigma | 0.86 | (0.76, 0.96) |
| Alpha | 3.09 | (2.31, 3.93) |

Table 2: Results from Dorsal Model and Ventral Model 2, where brightness is predicted by temperature, precipitation and habitat type.
| Group-Level Effects | Estimate | 95% CI |
| --- | --- | --- |
| Phylogenetic Error (sd) | 0.16 | (0.01, .039) |

We used the ELPD scores of the WAIC analysis to compare Models 1 and 2 to the null. We found that both predicted dorsal plumage brightness better than the null model (Table 3). However, the standard error of the ELPD scores of Models 1 and 2 overlapped, so that we are unable to conclude which of these two models better predicted dorsal plumage brightness. For the ventral plumage, Models 1 and 2 also better predicted ventral brightness than the null model, but in this case Model 2, which included habitat, was a better predictor of ventral plumage brightness than Model 1, which did not include habitat.

**Table 3:**
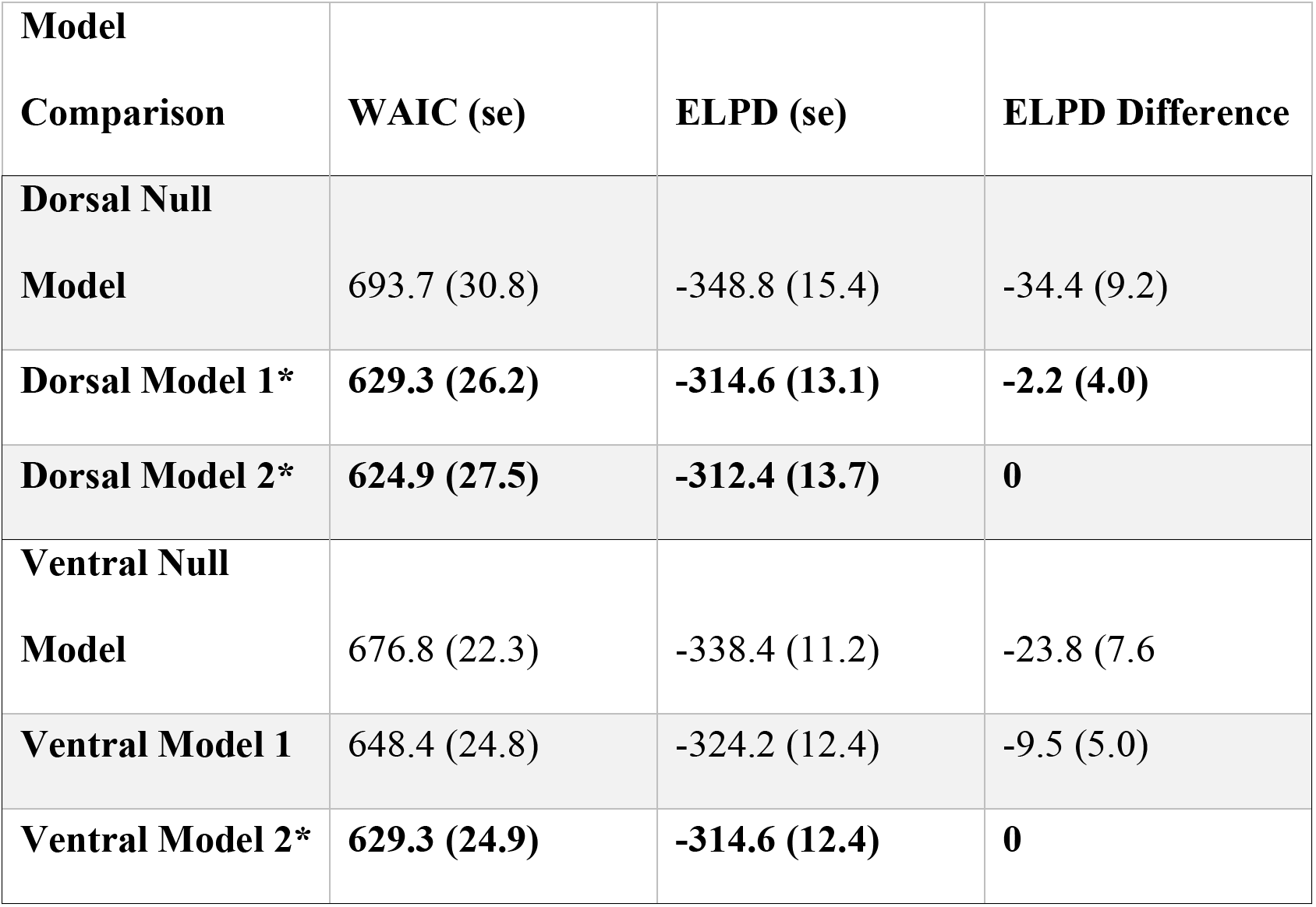
Model comparison for Dorsal Models and Ventral Models. The first column shows the Widely Applicable Information Criterion (WAIC) score with the standard error of the estimate in parentheses. The second model shows the Estimated Log-Predictive Density (ELPD), or the difference in the model’s predictive accuracy, with standard error in parentheses. The third column provides the difference between the ELPD scores and the model with the highest predictive accuracy. In Model 1, brightness is predicted by temperature, precipittaion and their interaction. In Model 2, brightness is predicted by temperature, precipitation and habitat type. For both venter and dorsum, Models 2 (in bold) have the highest predictive accuracy for the given data, and the null models have the lowest predictive accuracy. In the Dorsal models, the standard error of the ELPD difference for Models 1 and 2 overlaps, which means we cannot determine which of the two models has the highest ELPD score.

## Discussion

Gloger’s rule is a longstanding ecogeographic principle predicting that birds and mammals that inhabit rainier and warmer climates tend to have darker plumage and pelage color than their counterparts (intra-as well as interspecific) from drier and cooler places (Gloger 1833, Rensch 1929, Mayr 1942, 1963, Delhey 2017, 2019). Here, we found strong support in the Furnariidae for the predicted relationship between brightness and precipitation. In contrast, we found that furnariid species tended to be darker in cooler climates, contrary to the second prediction of Gloger’s rule, but consistent with a pattern dubbed Bogert’s rule or thermal melanism, which is often observed in ectothermic animals (Clusella-Trullas et al. 2018). We also found a credible positive interaction between precipitation and brightness, meaning that the negative relationship between precipitation and plumage brightness becomes stronger in cooler climates (Figures 1 and 2). Finally, forest-based lineages tended to have darker plumage than nonforest-based lineages (Figure 3), consistent with a previous study on furnariids and other closely-related families (Marcondes and Brumfield 2019). But, here, we expanded on that previous finding by showing that that tendency for birds to have darker plumage in darker habitats persists even after accounting for the effects of climate. This indicates that climate and light environments have separate but complementary effects in driving macroevolutionary patterns of plumage color variation in birds.

### Gloger’s rule, precipitation and temperature

Gloger (1833) wrote that “melanins […] increase with higher temperature and humidity” (translation from the German from Delhey 2019), implicating both climatic variables in the rule that would become his namesake. But Rensch (1936), in the first major discussion of Gloger’s rule in English, downplayed the role of temperature, placing more importance on humidity (reviewed by Delhey 2019).

The test of time—and of modern quantitative techniques—have validated Rensch’s (1936) emphasis on humidity. Intra-(e.g., Rising et al. 2009, Amar et al. 2014, Marcondes et al. in review) and interspecific (e.g., Delhey 2018, Delhey et al. 2019) comparisons, including this study, have consistently failed to find support for a tendency for birds to be darker in warmer places. Our Models 1 and 2 showed a positive effect of temperature on brightness, particularly in rainy climates (see below). This is diametrically opposite to Gloger’s (1833) formulation, but in accordance with intraspecific findings in the Black Sparrowhawk *Accipiter melanogaster* (Amar et al. 2014), Savannah Sparrow *Passerculus sandwichensis* (Rising et al. 2009) and Variable Antshrike *Thamnophilus caerulescens* (Marcondes et al. in review), as well as comparative results from analyses of the Australian avifauna (Delhey 2018) and the world’s passerines (Delhey et al. 2019). These findings are consistent with Bogert’s rule, a lesser known ecogeographical rule usually considered to apply only to ectothermic animals (Clusella-Trullas et al. 2018, Delhey 2018, 2019). This rule predicts animals should be darker in cooler climates to enhance thermoregulation. The consistency of results showing the same pattern in birds suggests that Bogert’s rule may be applicable to endothermic animals as well, although we lack an understanding of its mechanistic underpinnings in that case. Experimental work would be better suited to advance our knowledge in that regard (Delhey 2018).

Our models showed a strong interaction between precipitation and temperature (Figures 1 and 2). In cooler temperatures, the correlation between greater precipitation and lower brightness was stronger than in warmer temperatures. For illustration, consider four species of furnariids, each occupying a different climatic regime (Figure 2): the Peruvian Recurvebill *Syndactyla ucayalae* (warm/rainy), the Necklaced Spinetail *Synallaxis stictothorax* (warm/dry), the Itatiaia Spinetail *Asthenes moreirae* (cool/rainy), and the Cream-rumped Miner *Geositta isabellina* (cool/dry). Our results suggest that the two species inhabiting dry climates are expected to be brighter than the two species inhabiting rainy climates. But the difference in brightness between the species inhabiting a cool/dry and a cool/rainy climate should be greater than the difference in brightness between the species inhabiting a warm/dry and a warm/rainy climate. This is indeed what our data show. The difference in the first principal component of dorsal brightness between *Geositta isabellina* (cool/dry) and *Asthenes moreirae* (cool/rainy) was 0.3328, whereas the difference in the first principal component of dorsal brightness between *Synallaxis stictothorax* (warm/dry) and *Syndactyla ucayalae* (warm/rainy) was 0.1544.

These results can be contrasted with those of Delhey et al. (2019), who, like us, found support for Gloger’s rule for precipitation and Bogert’s rule for temperature across the world’s passerines, but did not test for their interaction. Delhey et al. (2019) proposed a general framework whereby the effect of temperature on plumage brightness has a quadratic shape, with birds being brighter at low and high temperatures and darker in intermediate temperatures, given the same levels of precipitation. Due to the credible interaction effect we found, our results do not conform to that framework. Instead, they suggest a more nuanced scenario: birds are lighter in cool and dry climates, especially for the ventral plumage, but in cool and rainier climates the effect of precipitation becomes more prevalent, leading to darker plumage (Figure 2). The difference between ours and Delhey et al.’s (2019) conclusions highlights how findings at a more broadly inclusive level (all passerines) may not be directly translatable to a more restricted clade (Furnariidae). This may be because the furnariids include proportionally fewer species occupying very cold climates relative to the passerines as a whole. The minimum temperature in our dataset was 1.7°C, whereas in Delhey et al.’s dataset it was lower than −10°C. Those species from very cold climates, which are also usually dry climates, could be driving the results observed when considering all passerines.

### Gloger’s rule, precipitation and habitat type

Numerous studies have shown that bird species of dark light environments (*e.g*. forests) tend to be darker than their relatives from open habitats, a pattern attributed to natural selection for crypsis (Endler 1993, McNaught and Owens 2002, Gomez and Thery 2004, Dunn et al. 2015, Maia et al. 2016, Shultz and Burns 2017, Marcondes and Brumfield 2019), but these studies have been conducted largely separately from investigations of Gloger’s rule (*e.g*., Delhey 2018, Delhey et al. 2019). We used a model with temperature, precipitation and habitat type as predictors of brightness to calculate contrasts between the posterior distributions of the phylogenetic mean of brightness in each habitat, while controlling for differences in the climatic variables. These contrasts showed that species from bright light environments (nonforest) are credibly brighter, ventrally, than those from intermediate light environments (forest edge and canopy), followed by species occupying the forest interior (Figure 3). Results were similar for the dorsal plumage, except that there was no difference between nonforest and intermediate habitats (Figure 3).

Zink and Remsen (1986) suggested background matching as the main adaptive mechanism responsible for Gloger’s rule. The aforementioned comparative work and our results corroborate this. Birds tend to be darker in darker (forested) habitats. Because forest habitats also tend to receive more precipitation (for example, precipitation in our dataset, mean±sd: forest species, 2009±611 mm/year; intermediate habitat, 1597±700 mm/year; nonforest 852±631 mm/year), the correlation between brightness and habitat could be spuriously driven by climate. Our results show that is not the case. The difference in brightness across habitats persists even after controlling for climatic variables, demonstrating that they have separate effects on the evolution of plumage brightness.

Zink and Remsen (1986) also suggested that “humidity per se presumably has little direct influence”. Because our Model 2 showed negative correlations between brightness and precipitation, even while including habitat as a predictor, we disagree. Higher precipitation, by itself, does correlate with darker birds. A potential explanation for this is protection against feather-degrading bacteria. It is well-documented that increased melanization makes feathers more resistant to feather-degrading bacteria (Goldstein et al. 2004, Gunderson et al. 2008), and that these bacteria are common on plumages of wild birds (Burtt and Ichida 1999, 2004, Kent and Burtt 2016). However, before it can be conclusively said that feather-degrading bacteria drive increased pigmentation in birds living in rainier habitats, evidence is needed that these bacteria are in fact more abundant in more rainy habitats.

### Gloger’s rule and vegetation density

Delhey (2018) used remote sensing data to show that, in Australia, birds tend to be darker in more heavily-vegetated areas. This is similar to, and consistent with, our findings. But our analyses based on habitat preference offer further insight, because bird species occupy habitat types differentially even within the same locality, a pattern that cannot be captured by remote sensing-based metrics of vegetation cover. For example, at a typically used resolution, remote sensing data may show that a 30 m x 30 m cell is covered in very dense, tall vegetation (rainforest). But different species of furnariids occupying that cell may experience diverse light environments. For example, in western Amazonia that cell may be occupied by the Orange-fronted Plushcrown *Metopothrix aurantiaca* in the intensely sun-lit forest canopy and the Tawny-throated Leaftosser *Sclerurus mexicanus* in undergrowth vegetation near the forest floor in the dim forest interior.

Remote sensing analyses may also be complicated by the fact they are often based on museum specimens collected up to a few decades ago, before recent intense anthropogenic landscape change that will be reflected in remote sensing data. The landscape where a bird was collected many years ago may have little resemblance to the landscape at the same locality today.

Nevertheless, vegetation density by itself might also favor increased pigmentation, because greater melanin content makes feathers harder and more resistant to abrasion (Barrowclough and Sibley 1980, Burtt 1986, Bonser 1995). This is often considered in the context of abrasion from airborne particles, but it is conceivable that abrasion from vegetation might also be an important selective factor favoring heavier plumage pigmentation (Kale 1966, Burtt 1986, Surmacki et al. 2011, Kroodsma and Verner 2013), although this demands further empirical study.

## Conclusion

Gloger’s rule is a classic ecogeographic principle predicting animals should be darker in wetter and warmer regions. We have shown, based on comparative analyses of the Furnariidae, a family of >200 Neotropical passerine species, that the prediction related to precipitation is borne out in our data, but the prediction related to temperature is not. In fact, we found that furnariids tend to be darker in cooler regions. We also found a previously undescribed credible interaction of precipitation and temperature, whereby the negative effect of precipitation on plumage brightness becomes stronger under cool temperatures. Furthermore, we also showed that species in this family tend to be darker in darker light environments and that this effect persists even after controlling for the effects of climate.

Based on ours and previous results, we suggest that the pattern encapsulated by Gloger’s rule is produced by a combination of the partially correlated effects of habitat type, precipitation, and vegetation density. The effect of habitat type is driven by natural selection for enhanced crypsis in darker light environments (Zink and Remsen 1986, Endler 1993, McNaught and Owens 2002, Gomez and Thery 2004, Dunn et al. 2015, Maia et al. 2016, Shultz and Burns 2017, Marcondes and Brumfield 2019), whereas the effect of precipitation may be due to featherdegrading bacteria (Burtt and Ichida 1999, 2004, Goldstein et al. 2004, Gunderson et al. 2008, Kent and Burtt 2016), and the effect of vegetation density may be related to feather abrasion (Kale 1966, Burtt 1986, Surmacki et al. 2011, Kroodsma and Verner 2013), though the latter two effects still demand further empirical work to be conclusively demonstrated. It is also still unclear how the effects of temperature fit into this scenario.

## Data accessibility statement

Color data is deposited on Dryad under DOI, 10.5061/dryad.s86434s. Climatic data will be deposited on Dryad upon acceptance for publication.

## Acknowledgments

This work was partially supported by NSF grant DEB-1146265 to RTB and by a “Science Without Borders” doctoral fellowship from Brazil’s National Council for Scientific and Technological Development to RSM (CNPq; 201234/2014–9).

